# Characterisation of palytoxin from an undescribed Palythoa (Anthozoa: Zoantharia: Sphenopidae) with significant in vitro cytotoxic effects on cancer cells at picomolar doses

**DOI:** 10.1101/292219

**Authors:** Ludovic Sawelew, Frédéric Gault, Christopher Nuccio, Yvan Perez, Jean Lorquin

## Abstract

Palytoxin (PlTX), a large polyhydroxylated compound, is among the most potent non-peptide toxin in marine organisms known so far. The literature emphasizes the sodium/potassium pump (NaK) as the privileged target for PlTX when exerting its toxic effects. In this study, we focused on an undescribed species (*Palythoa* sp. Pc001), a coral species belonging to the genus *Palythoa* routinely cultivated in aquariums. We demonstrated that this species contains one of the highest yields of pure PlTX production ever found, 2.22 ± 0.41 mg PlTX per gram of wet *Palythoa*. Using molecular data combined with external morphology, we identified *Palythoa* sp. Pc001 as the sister species to *Palythoa* aff. *clavata*. Further, the clade of a symbiotic *Symbiodinium* sp. was characterised by DNA barcoding and pigment content. Molecular data showed that *Palythoa* sp. Pc001 contains ‘generalist’ *Symbiodinium* belonging to clade C. This paper also describes for the first time the localisation of PlTX and *Symbiodinium* cells in tissues of a highly toxic *Palythoa* species. PlTX toxicity was assayed on 72 h-cultured murine and human cancer cells *versus* the normal human dermal fibroblast (NHDF; PC C12300) cell line. Using MTT colorimetric assay and quantitative videomicroscopy, our results showed much higher *in vitro* cytotoxic activity on cancer cells (IC_50_ 0.54 ± 0.05 × 10^−12^ M) than on non-cancerous ones (IC_50_ > 1 × 10^−6^ M). Such a strong differential effect has never been reported with respect to the most potent NaK ligands (cardiac glycosides) described so far. Moreover, PlTX displayed similar *in vitro* growth inhibitory activity in rodent and human cancer cells, although the NaK in rodents displays a double mutation in the α1-subunit that usually decreases the sensitivity to others cardiac glycosides like ouabain, when compared to human cells. This work demonstrates, first, that picomolar concentrations of PlTX have significant higher cytotoxic effects on cancer cells than on non-cancerous ones, and secondly, that this *in vitro* antitumor effect would not be entirely relied onto its canonical targeting to the NaK α-subunit. Thus, PlTX ranks amongst highly potent anti-cancer drugs as it targets cancers while potentially minimizing the drug’s side effects on healthy cells.

## Introduction

Many organisms such as animals, plants and bacteria are known to secrete poisonous substances in their environment to protect themselves from aggressors. These toxins may also be very useful in cancer therapy due to their high cell-killing potency [1–3]. PlTX, one of the most toxic natural compounds known to date, is a nonprotein marine toxin which consists of a long, partially unsaturated (with eight double bonds) aliphatic backbone with spaced cyclic ethers and 64 chiral centers [4]. Initially isolated from a *Palythoa* species (Anthozoa: Zoantharia: Sphenopidae) [5], it can also be found in numerous other marine organisms from the same ecological region [6]. Moreover, several analogues of PlTX were discovered in various organisms (Table 1). Currently, the biogenetic origin of PlTX in *Palythoa* remains unclear. The leading hypothesis is that the toxin is synthesised by symbiotic dinoflagellates [7, 8] and/or by bacteria [9, 10]. To date*, Palythoa heliodiscus* is the largest PlTX producer known among zoanthids, providing high yield of PlTX (1 mg/g wet *Palythoa*) and deoxy-PlTX (3.51 mg/g wet *Palythoa*) [11]. However, in most cases the toxin yield is very low. As a consequence, and because of sanitary problems due to toxic *Ostreopsis* spp. outbreaks [12], there is a need for high-yield sources of PlTX so as to facilitate its functional characterisation. Sales from aquarium trade of zoanthids containing PlTX have also caused domestic intoxications after inhalation of steam [13, 14], dermal contact [15] or water spray in the eyes [16, 17].

**Table 1.**
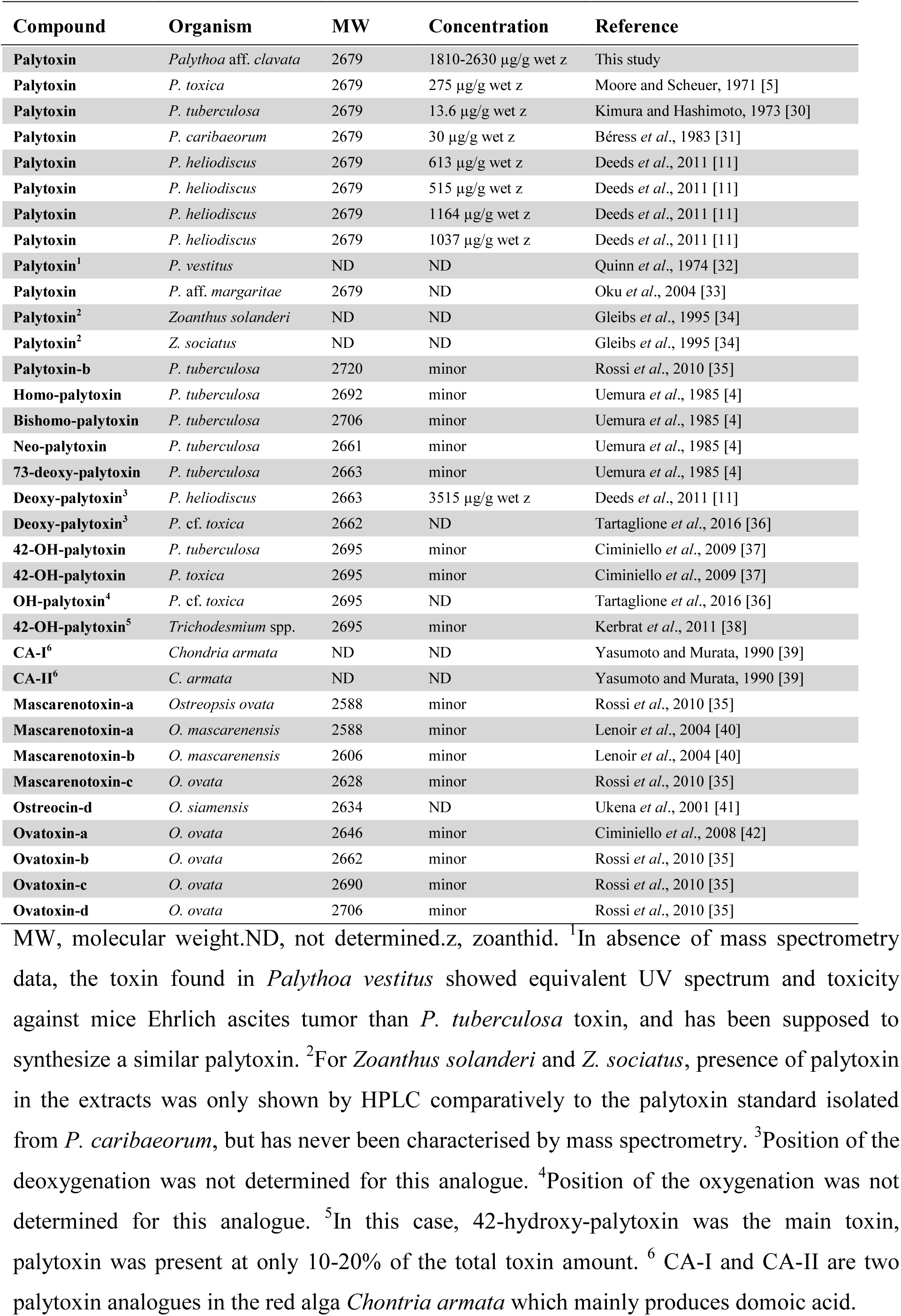
Summary of the known molecules in the palytoxin family in Zoantharia, Cyanobacteria, algae and dinoflagellates.

PlTX and its analogues bind and transform the Na^+^/K^+^-ATPase pump (NaK) into an open channel. The PlTX binding site is coupled to other cardiac glycosides (CGs), such as ouabain which is known to bind the α-subunit of NaK [18]. NaK α-subunit isoforms have a similar affinity for all CGs [19]. However, recent kinetic experiments revealed that PlTX binding dissociation resulting from ouabain competition was incomplete, suggesting an additional, ouabain-insensitive, PlTX binding site [2]. The ensuing ionic passive transport through NaK leads to the depolarization of the plasma membrane [20, 21] and affects the mechanisms controlling intracellular calcium (Ca^2+^) concentration [22]. The intracellular Ca^2+^ increase is related to a long-lasting and gadolinium-sensitive Ca^2+^ influx suggesting a possible involvement of stretch-activated channels in PlTX-induced cytotoxicity [23]. Furthermore, PlTX mediates disruption of the actin cytoskeleton and cytomorphological changes [24, 25]. The massive increase of Ca^2+^ in the cytosol triggers the activation of cellular signalling pathways. As is the case with various CGs [26], mitogen activated protein kinases (MAPKs) mediate PlTX-stimulated signalling and relay a variety of signals to the cellular machinery that regulates cell fate and function [27].

In this study, we have shown that aquacultured zoanthids from an undescribed species belonging to the *Palythoa* aff. *clavata/sakurajimensis* complex represent a new PlTX source with the highest yield ever found. While PlTX has been primarily identified as a tumour promoter when combined to carcinogenic compounds [28, 29], the PlTX isolated from *Palythoa* sp. Pc001 exhibited high *in vitro* cytotoxic effects on cancer cells at picomolar doses. The IC_50_ concentrations calculated from MTT colorimetric assay associated with quantitative videomicroscopy showed that *in vitro* cytotoxicity is over 1000 times higher toward cancer cells than non-cancerous NHDF cells. Moreover, comparative MTT colorimetric assay for *in vitro* growth inhibitory effects on murine *versus* human cancer cell lines suggested that the targeting of the NaK α-subunit is not the sole mechanism underlying the cytotoxic effects of PlTX. Based on these results PlTX may be a very promising anti-cancer agent.

MW, molecular weight.ND, not determined.z, zoanthid. ^1^In absence of mass spectrometry data, the toxin found in *Palythoa vestitus* showed equivalent UV spectrum and toxicity against mice Ehrlich ascites tumor than *P. tuberculosa* toxin, and has been supposed to synthesize a similar palytoxin. ^2^For *Zoanthus solanderi* and *Z. sociatus*, presence of palytoxin in the extracts was only shown by HPLC comparatively to the palytoxin standard isolated from *P. caribaeorum*, but has never been characterised by mass spectrometry. ^3^Position of the deoxygenation was not determined for this analogue. ^4^Position of the oxygenation was not determined for this analogue. ^5^In this case, 42-hydroxy-palytoxin was the main toxin, palytoxin was present at only 10-20% of the total toxin amount. ^6^ CA-I and CA-II are two palytoxin analogues in the red alga *Chontria armata* which mainly produces domoic acid.

## Materials and methods

### *Palythoa* sp. Pc001 origin

*Palythoa* sp. Pc001 specimens used in this study were provided from the aquarium trade industry (Indonesia) and bred at Coral Biome’s facility (Marseille). Specimens were maintained in a closed system aquaculture connected to a biological filter fed by artificial seawater (Instant Ocean salts, from Seachem, Madison, USA). Specimens were maintained at 26 ± 1°C and exposed to a daytime photoperiod of 12 h with an irradiance of 70 mol.quanta m^−2^.s^−1^and fed daily with fish food pellets (Formula One, from Ocean Nutrition, Essen, Belgium) to maximize growth rate.

### DNA extraction, PCR amplification and sequencing

To amplify both *Palythoa* and *Symbiodinium* genes, tissue sample consisting of five tentacles joined by a small piece of polyp oral disc was placed in 80% alcohol. Total genomic DNA was extracted using the DNAeasy Kit (Qiagen, Valencia, CA). Two commonly used DNA barcode markers were amplified to identify the *Palythoa* sp. Pc001 specimens: the mitochondrial cytochrome oxidase subunit I (COI) and the nuclear internal transcribed spacer region of ribosomal DNA (ITS-rDNA, internal transcribed spacer 1, 5.8S ribosomal RNA gene, internal transcribed spacer 2). An ITS-rDNA sequence of approximately 750-850 base pairs was amplified using the primers Zoanf-ITS (5’ - CTT GAT CAT TTA GAG GGA GT - 3’) and Zoanr-ITS (5’ - CGG AGA TTT CAA ATT TGA GCT - 3’). The PCR program was carried out as follow: an initial denaturing step at 94°C for 3 min, followed by 35 cycles of 1 min denaturation at 94°C, 1 min annealing at 50°C, 2 min extension at 72°C followed by 10 min at 72°C. The portion of the mitochondrial COI gene of approximately 650 base pairs was amplified with the following primers: HCO2198 (5’ - TAA ACT TCA GGG TGA CCA AAA AAT CA - 3’) and LCO1490 (5’ - GGT CAA CAA ATC ATA AAG ATA TTG G - 3’). PCR amplification was performed as follow: an initial denaturing step at 94°C for 2 min followed by 5 cycles of 15 s denaturation at 92°C, then 45 s annealing at 48°C followed by an incremental increase until 72°C in 1 min, then 1 min 30 extension at 72°C, followed by 30 cycles of 15 s denaturation at 92°C, 45 s annealing at 52°C, 45 s extension at 72°C, followed by 7 min at 72°C.

To identify the *Symbiodinium* clade associated to *Palythoa* sp. Pc001, an ITS2-rDNA sequence (5.8S ribosomal RNA, partial sequence; internal transcribed spacer 2, complete sequence; 28S ribosomal RNA gene, partial sequence) of approximately 250-300 base pairs was amplified using the specific primers ITS2-F1 (5’ - GAA TTG CAG AAC TCC GTG - 3’) and ITS2-R2 (5’ - ATA TGC TTA AAT TCA GCG GGT - 3’). PCR amplification was performed under stringent conditions to specifically target the *Symbiodinium* genes instead of coral ones: an initial denaturing step at 94°C for 3 min followed by 12 cycles of 45 s denaturation at 94°C, 45 s annealing at 58°C and an incremental decrease of 0.5°C every cycle, 1 min extension at 72°C, followed by 20 cycles of 45 s denaturation at 94°C, 45 s annealing at 52°C and 1 min extension at 72°C followed by 7 min at 72°C.

After amplification, all PCR fragments were visualised by denaturing gradient gel electrophoresis and were sequenced in both directions using the amplicon primers with an ABI 96-capillary 3730XL sequencer at Eurofins genomics (Ebersberg, Germany).

### Phylogenetic analyses

Two data sets were used for molecular analyses: dataset 1 for *Palythoa* species with ITS-rDNA and COI concatenated sequences and dataset 2 for *Symbiodinium* with ITS2-rRNA sequences. All sequences obtained in this study were first checked using NCBI BLAST, then aligned with orthologous sequences available in public databases using CLUSTALW implemented in BioEdit v7.1.9 (http://www.mbio.ncsu.edu/bioedit/bioedit.html). The program MUSCLE [43] was also used to carry out a multiple alignment based on the *Symbiodinium* ITS2-rDNA sequences. The MODELTEST v3.0b4 program [44] was used to identify the best model of DNA evolution using Bayesian information criterion (BIC). Molecular analyses were conducted through neighbour-joining (NJ) and maximum likelihood (ML) methods, using MEGA v6.0 [45] and Bayesian inference (BI) employing MrBayes v3.2 [46]. Topological robustness was determined using 100 non-parametric bootstrap replicates for NJ and ML analyses. For BI, Markov Chain Monte Carlo searches were done with four chains for 1,000,000 generations, with a random starting tree, default priors and Markov chains (with default heating values) sampled every 1,000 generations.

### MALDI-IMS analyses and localisation of *Symbiodinium* cells in *Palythoa* Pc001 tissues

Polyps were collected from cultured colonies in aquariums, harvested with care, and then quickly frozen in a container of isopentane plunged in liquid nitrogen. After 5 min, dry polyps were stored at −80°C overnight. Tissue sections were cut using a Leica CM 1900 UV Microsystems cryostat (Leica Microsystems SAS) with a microtome chamber and a specimen holder chilled at −20°C. Sections of 18 µm were made in crosswise directions at three levels of the polyp body. Lengthwise sections were also made in some specimens. Sections were thaw mounted onto Indium Tin Oxide coated microscopic slides (Bruker Daltonics) adapted for matrix-assisted laser desorption ionization imaging mass spectrometry (MALDI-IMS), and onto superfrost plus slides (Thermo Scientific) for epifluorescence imaging. Both types of target slides were dried in a desiccator for 45 min. Polyps sections were scanned before matrix deposition with a histology slide scanner (Opticlab H850 scanner, Plustek). Then, 2,5-dihydroxybenzoic (DHB) acid (Bruker, Daltonics), 30 mg/mL in 50/50 methanol (MeOH)/H_2_O 0.1% trifluoroacetic acid (TFA) was used as a matrix and applied on tissue sections using an automatic matrix sprayer (TM-Sprayer, HTX Technologies). MALDI calibration was carried out manually with a Peptide Calibration Standard 2 (Bruker Daltonics). MALDI-IMS analyses were performed using an Ultraflextreme mass spectrometer controlled by the FlexControl 3.3 software (Bruker Daltonics) in positive reflectron mode. MALDI-IMS sequences were created with FlexImaging 3.0 and the measurement region were manually defined with the previous histological images. The spatial resolution was set at 50 µm with a laser diameter of 20 µm, and 300 laser shots were accumulated for each spot. The laser power was optimised at the start directly on tissue and then fixed for the overall MALDI-IMS experiment. The images were opened with SCILS Lab v2.5 in RAW data with a baseline subtraction to maintain resolution of the average mass spectrum. Due to the high intensity of the PLTX signal, root mean square normalisation was chosen for a better visualization of the distribution. Endogenous auto fluorescence of chlorophyll and peridinin pigments of the *Symbiodinium* cell was revealed under an inverted microscope (Nikon Eclipse TE2000-U) using a filter cube B-2E/C corresponding to a medium band blue excitation. Images were acquired with NIS-Elements BR v2.30 imaging software.

### PlTX purification

About one gram of fresh *Palythoa* sp. Pc001 was gently detached from the substrate with a scalpel, chopped into several pieces and placed in 20 mL of 80% (v/v) MeOH in milliQ H_2_O. After agitation for 12 h at 4°C, the extract was centrifuged (8,000 g, 20 min), the pellet rinsed with 5 mL milliQ H_2_O, centrifuged (8,000 g, 20 min) and the supernatants were pooled. MeOH was evaporated using a rotavapor and the aqueous phase extracted several times with dichloromethane to remove carotenoid and chlorophyll pigments. The red-color (RC) organic phases were pooled, evaporated to dryness and kept at 4°C in darkness for further pigment identification. The remaining aqueous phase that contained PlTX was evaporated and deposited onto a two-centimeter diameter glass column filled with 10 cm^3^ of C_18_ reversed phase powder (Lichroprep RP_18_, from MERCK, France). The column was washed with acidified H_2_O (0.2% (v/v) formic acid), then with 50% (v/v) MeOH in acidified H_2_O. PlTX was finally eluted with 75% (v/v) MeOH in acidified H_2_O, and a dried pale-yellow solid of pure PlTX was obtained by N_2_ flow evaporation. The toxin was solubilised in dimethylsulfoxide (DMSO) and quantified by using high performance liquid chromatography (HPLC, see below). Routinely, preparations containing 100 µg PlTX in 100 µL DMSO were stored at 4°C for up to 6 months. From the last purification step, a yellow carotenoid (YC) fraction retained on the Lichroprep-RP column was eluted by pure MeOH, evaporated to dryness and stored at 4°C in darkness for further identification.

### HPLC analyses

Solvents were of HPLC grade and obtained from Biosolve (Dieuze, France). To control the purity of the PlTX fraction and quantify the toxin, 2-5 µg of the sample in milliQ H_2_O were injected and analysed by reverse-phase (RP) HPLC with a Waters equipment composed of a 1525 binary pump, a 2996 diode array detector, a 7725i Rheodyne injector fitted with a 20-µL loop, and a temperature control system. Files were acquired by the Empower software. Separations were carried out on a Waters Symmetry C_18_ column (4.6 × 100 mm, ODS2, 5 µm) protected with a guard cartridge. The elution was performed at 30°C at a flow rate of 0.8 mL/min and using a linear gradient of MeOH (eluent A) in acidified H_2_O with 0.2 % (v/v) acetic acid (eluent B), from 5 to 100 % A during 20 min. PlTX was visualised at 263 nm and total spectra were analysed from 200 to 700 nm by the software to control the purity. Commercial PlTX (from WAKO Pure Chemical Industries, Japan) was redissolved in DMSO and used for calibration. Data were averaged from three injections. The yield of PlTX in *Palythoa* sp. Pc001 (expressed as mg/g of wet *Palythoa*) was determined by averaging the yields calculated from 12 animal extractions. Pigments from *Symbiodinium* and *Palythoa* sp. Pc001 (RC and YC) extracts were also analysed by the same HPLC device but using an Agilent Technologies Zorbax Eclipse XDB-C_8_ column (4.6 × 150 mm) eluted with the gradient method of Zapata *et al*. [47].

### Mass spectrometry

Matrix-assisted laser desorption ionization time-of-flight (MALDI-ToF) analyses were performed on a Microflex II mass spectrometer (Bruker, Germany). A 10 mg/mL solution of 2,5-DHB in 70/30 acetonitrile/H_2_O 0.1 % TFA was used as matrix [48]. From a solution of 1 mg/mL of purified toxin in milliQ H_2_O, 0.8 µL were mixed with 0.8 µL of the matrix solution, deposited and allowed to dry at room temperature. Data were acquired in a positive reflectron mode, the range was set from 600 to 5000 Da, and pulsed ion extraction was fixed at 150 ns. Ions formed upon irradiation by a smartbeam laser using a frequency of 200 Hz. The laser irradiance was set to 45-50% (relative scale 0-100) arbitrary units according to the corresponding threshold required for the applied matrix system. Mass spectra were treated with the Flex Analysis software (Bruker, Germany) and no smoothing or baseline subtraction was performed. Purified PlTX was also analysed by ESI-MS/MS on a Q-ToF Synapt G1 High Definition mass spectrometer (Waters, UK) mounted with a nanospray ionization source and a V-mono-reflectron, in the positive mode. The following source settings were used: capillary voltage 3.2 kV, sampling cone 40 V, extraction cone 4 V, source temperature 120°C, desolvation gas flow 200 L/h (N_2_) at a temperature of 150°C, and trap collision energy 38 eV. The calibration was performed with a solution of CsI 1 mg/mL and used in the 100-3200 mass range with a precision of +/- 3 ppm. A solution of purified PlTX (1 mg/mL) in MeOH/H_2_O (1:1) was diluted 25 times (15 µM final concentration) and injected via a syringe pump in the nano-source at a flow rate of 3 µL/min.

### Carotenoids and chlorophylls identification

RC and YC fractions obtained previously as dried matter (see PlTX purification) were each dissolved in 1 mL of MeOH and 10-20 µL were directly analysed by HPLC on a C_8_ reverse-phase column. For comparison, photosynthetic pigments were also extracted from a pellet containing fresh *Symbiodinium* cells with 5 mL of a MeOH-H_2_O (4:1) mixture under agitation with a magnetic barrel for 2 h at 4°C in darkness. After centrifugation (10,000 g, 5 min), the coloured supernatant was slightly evaporated and 20 µL were directly analysed by HPLC. *Symbiodinium* cells were isolated according to a protocol adapted from Perez and Weis [49]. This involved cutting in half three polyps and homogenizing each half in a glass tissue grinder with 1 mL of filtered artificial seawater. After decantation and leaving pieces of *Palythoa* sp. Pc001 in the bottom grinder, the six solutions were pooled and centrifuged at 450 g for 5 min to pellet the algae which were then washed in three changes of seawater.

### Cell lines and culture media

All medium components were purchased from Lonza (Westburg, Germany). The cell lines were purchased from the following bank collections: the American Type Culture Collection (ATCC, Manassas, USA); the European Collection of Cell Cultures (ECACC, Salisbury, UK); the Deutsche Sammlung von Mikroorganismen und Zellkulturen (DSMZ, Braunschweig, Germany); the Cell Line Services (CLS, Eppelheim, Germany); PromoCell (PC, Europe). The non-cancerous cell line used was the normal human dermal fibroblast (NHDF; PC C12300). The cancer cell lines sensitive to pro-apoptotic stimuli used here included the human Hs683 oligodendroglioma (ATCC HTB-138) and the mouse B16F10 melanoma (ATCC CRL6475) cell lines [50, 51, 52]. The cancer cell lines displaying various levels of resistance to pro-apoptotic stimuli included the human A549 non-small-cell lung cancer (NSCLC; DSMZ ACC-107), the human U373 glioblastoma cells (ECACC08061901) [50, 53], and the rodent 9L gliosarcoma cells. The human HBL-100 (CLS 330178) epithelial cell line was also included as a non-cancerous but transformed model. All cell lines except NHDF were cultured in RPMI 1640 culture medium supplemented with 10% heat-inactivated FBS and a mixture of glutamine 0.6 mg/mL, penicillin 200 IU/mL, streptomycin 200 IU/mL and gentamycin 0.1 mg/mL (all at final concentration). The NHDF cell line was cultured in an MEM culture medium with 5% heat-inactivated FBS and antibiotics as described above.

### MTT colorimetric assay and quantitative videomicroscopy

The MTT colorimetric assay was used in order to determine the *in vitro* IC50 growth inhibitory concentrations of PlTX. The MTT colorimetric assay was performed on the panel of cell lines listed above as detailed in [53]. Cell lines were plated according to their growth rate and left to adhere and grow for 24 h before treatment. Cells were then treated for 72 h with different concentrations of purified PlTX in DMSO ranging from 1 pM to 1 µM, with a semi-log concentration increase. After the treatment period, cells were incubated with the 3-(4,5-dimethylthiazol-2-yl)-2,5-diphenyl tetrazolium bromide (MTT) solution to enable mitochondrial reduction. Blue formazan was solubilised with DMSO, and optical density was measured at 570 nm (reference wavelength 610 nm). The IC_50_ concentrations were calculated from 96-well plates in which each experimental condition was carried out in six replicates except for the control, which was carried out in 12 replicates.

Direct visualization of the PlTX-induced effects on the cell proliferation and morphology of human Hs683 and U373n glioma cells was performed by means of time-lapse computer-assisted phase contrast microscopy, *i.e*. quantitative videomicroscopy as previously described [54]. A picture of the same field was acquired every 4 min over a period of several hours. Movies were generated from these digitised images to enable a rapid viewing of the cell behaviour for the duration of the experiments in control versus treated experimental conditions. Experiments were carried out in tetraplicates in human Hs683 and U373 glioma cell lines in the absence (control) or the presence of PlTX at 0.01 and 1 nM.

## Results

### Morphological description and molecular typing of *Palythoa* sp. Pc001

The sand-encrusted button-like body, the oral disk and tentacles (approximately 60) are brown and punctuated with white dots (Fig 1). The oral disc, with a diameter of 10-15 mm, has green overtones with a whitish centre and a white strip that forms a radius crossing the centre of the oral disc. The sequences of ITS-rRNA and COI genes from 17 specimens of *Palythoa*(representing 10 species) and 4 specimens of *Zoanthus* (representing 4 species) were concatenated to increase the accuracy of the phylogenetic reconstructions. The total length of the concatenation based on the two genes was 1221 bp (770 and 451 pb ITS-rRNA and COI respectively). The best-fitting model K2+Γ was used for all methods. BI was performed with the following settings: the model employed two substitution types (nst = 2) with stationary state frequencies fixed to equal in order to get the K2 model (prset statefreqpr = fixed(equal)). Rate variation across sites was modelled using a gamma distribution (rates = gamma). Sequences from *Zoanthus kuroshio*, *Z. gigantus*, *Z. sansibaricus* and *Z. sociatus* were used as outgroups (Fig 1). The monophyly of *Palythoa* genus was highly supported whatever the method considered (100 bootstrap value (bv) for NJ /100 bv for ML /1 posterior probability (pp) for BI). Monophyletic lineages within the *Palythoa* genus were identified with good support values. The first clade (100/100/1) contained *Palythoa grandis* as the sister group to the *P. variabilis*/*P. heliodiscus* complex (100/97/1). The second clade consisted of *P. caribaeorum*, *P. tuberculosa* and *P. Mutuki*, although this group was moderately supported (76/73/1). *P. grandiflora* appeared as the sister species to this assemblage but with low bv values (66/69/1). The new sequences obtained from *Palythoa* sp. Pc001 grouped within a well-supported clade containing *P.* sp. *sakurajimensis* (Japan) and *P.* aff. *clavata* (Florida) (99/91/1). However, the sister group relationship between *Palythoa* sp. Pc001 and *P.* sp. *clavata* was supported by BI only (pp = 0.96).

**Figure 1.**
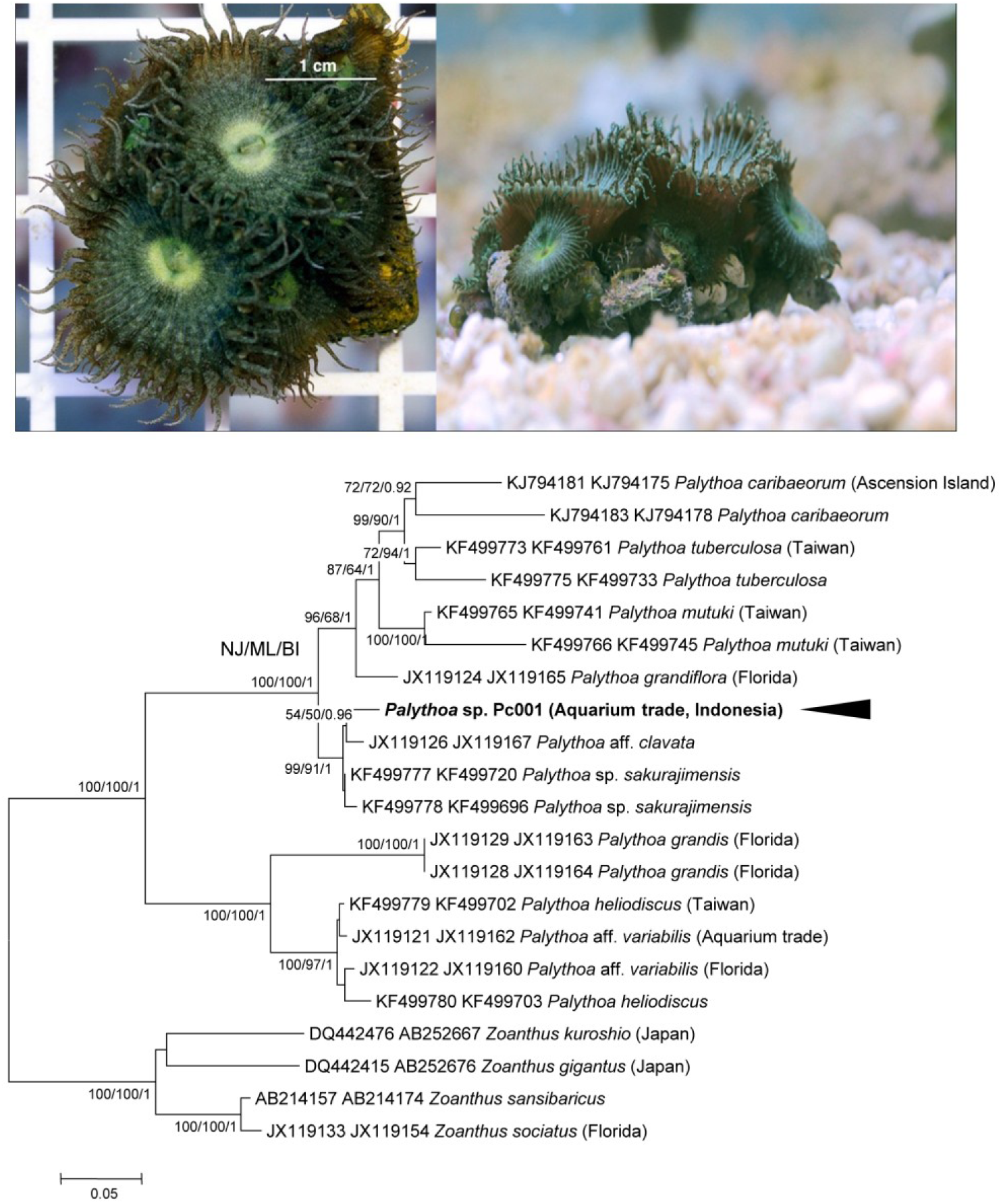
Morphotype and phylogenetical identification of *Palythoa* sp. Pc001 used in this study. Top: two colonies of *Palythoa* sp. Pc001 collected in Indonesia and routinely cultivated in aquariums at Coral Biome. Bottom: maximum likelihood tree calculated from a supermatrix including ITS-rRNA and COI sequences (1221 nucleotide positions) under the K2 + Γ model. The lnL value of this optimal tree is −5683.49. Support values obtained using different reconstruction approaches are indicated at nodes in the following order: neighbor joining (NJ), maximum likelihood (ML), and bayesian inference (BI). Support values are displayed when bootstrap values ≥ 70 or posterior probabilities p ≥ 0.90. Tree was rooted on the monophyletic assemblage consisting of 4 *Zoanthus* species. Black arrowhead indicates the sequence belonging to *Palythoa* sp. Pc001 used for palytoxin extraction. Sequences/species names from previous studies in regular font with GenBank Accession Numbers.

### Molecular typing of the *Symbiodinium* clade and pigment characterisation

After the sequencing step, the ITS2-rDNA sequence displayed clear chromatograms in both forward and reverse directions with no ‘double-peaks’, and therefore no cloning step was performed to investigate the intragenomic variability. The best fit model for data set 2 was K2+I. Phylogenetic analysis of ITS2-rDNA sequences, which were rooted on midpoint, yielded well-resolved trees in which the main clades previously described in cnidarians received high support values whatever the method considered (Fig 2). *Symbiodinium* ITS2-rDNA sequence isolated from the *Palythoa* sp. Pc001 grouped within the ‘generalist’ clade C (93/100/0.94). To check the origin of the chlorophylls and carotenoids, total pigments were extracted from purified *Symbiodinium* cells and analysed by RP-HPLC-DAD on a C_8_ column. The eluted pigments were identified as peridinin (major pigment, retention time 13.4 min, »max 474 nm) and its *cis*-isomer (minor, 13.8 min, 459 nm), fucoxanthin (major, 18.2 min, 468 nm) and its isomer (minor, 17.9 min, 461 nm), diadinoxanthine (major, 23.0 min, [422,446,477] nm) and dinoxanthin (minor, 24.0 min, [416,441,469] nm), chlorophyll *a* (major, 34.0 min, [429,616,662] nm) and its epimer (major, 34.4 min, [430,618,662] nm), and the very weak compounds chlorophyll *c*_*2*_ (9.1 min, 450 nm) and phaeophytin *a* (hypothetic, 36.0 min, [409,607,660] nm) (profile not shown). No β-carotene was detected. In parallel, the dichloromethane (RC) and the C_18_ (YC) fractions obtained during the PlTX purification process were also analysed for their pigment content. Identical pigments comprising carotenoids and chlorophylls were found in the RC fraction, since carotenoids only were present at lower doses in the YC fraction (data not shown), indicating that the liquid-liquid extraction using dichloromethane removed the totality of the chlorophylls and a major part of the carotenoids which facilitated the subsequent purification steps.

**Figure 2.**
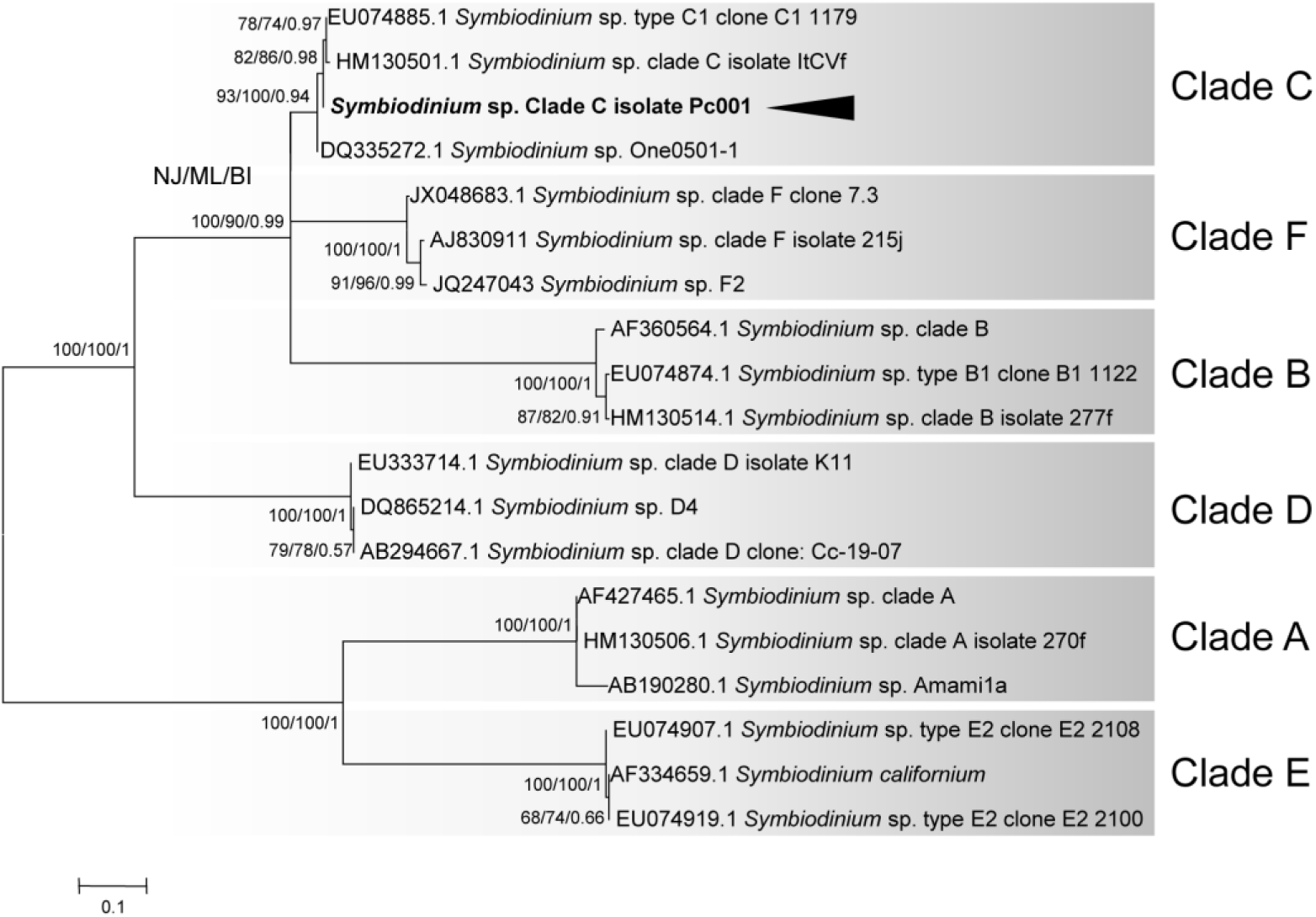
Phylogenetic identification of a *Symbiodinium* sp. from *Palythoa* sp. Pc001. Maximum likelihood tree calculated from ITS2-rRNA sequences (268 nucleotide positions) of *Symbiodinium* clades from various corals under the K2 + I model. The lnL value of this optimal tree is −1592.20. Support values obtained using different reconstruction approaches are indicated at nodes in the following order: neighbor joining (NJ) maximum likelihood (ML) bayesian inference (BI). Black arrowhead indicates the sequence belonging to the *Symbiodinium* clade identified in the *Palythoa* species used for palytoxin extraction. Sequences/species names from previous studies are indicated in regular font with GenBank Accession Numbers. On the right, *Symbiodinium* clades confirmed to be in symbiosis with various metazoans species (this study and previous studies) designated with shaded boxes.

### Purification, quantification and characterisation of PlTX

PlTX was eluted as a pure compound at 75% MeOH by HPLC and DAD acquisitions in the 200-700 nm range. The purified fraction was eluted as a symmetrical peak at about 80% MeOH on the C_18_-HPLC column (Fig 3A) by using a 0.2% acetic acid in the aqueous solvent (eluent B), the retention time was 14.8 min instead of 15.8 min in a 1% acid concentration. The purified fraction displayed two UV bands characteristic of PlTX at 233 and 263 nm. The purity was controlled by C_18_-HPLC-DAD in the entire UV-visible range, and then by C_8_-HPLC-DAD to verify the presence of residual carotenoid/chlorophyll pigments. No pigment or other compound was detected. Specimens of *Palythoa* sp. Pc001 were found to produce 2.22 ± 0.41 mg of PlTX/g wet *Palythoa (n = 11)*, as evaluated by HPLC analyses (Table 1).

**Figure 3.**
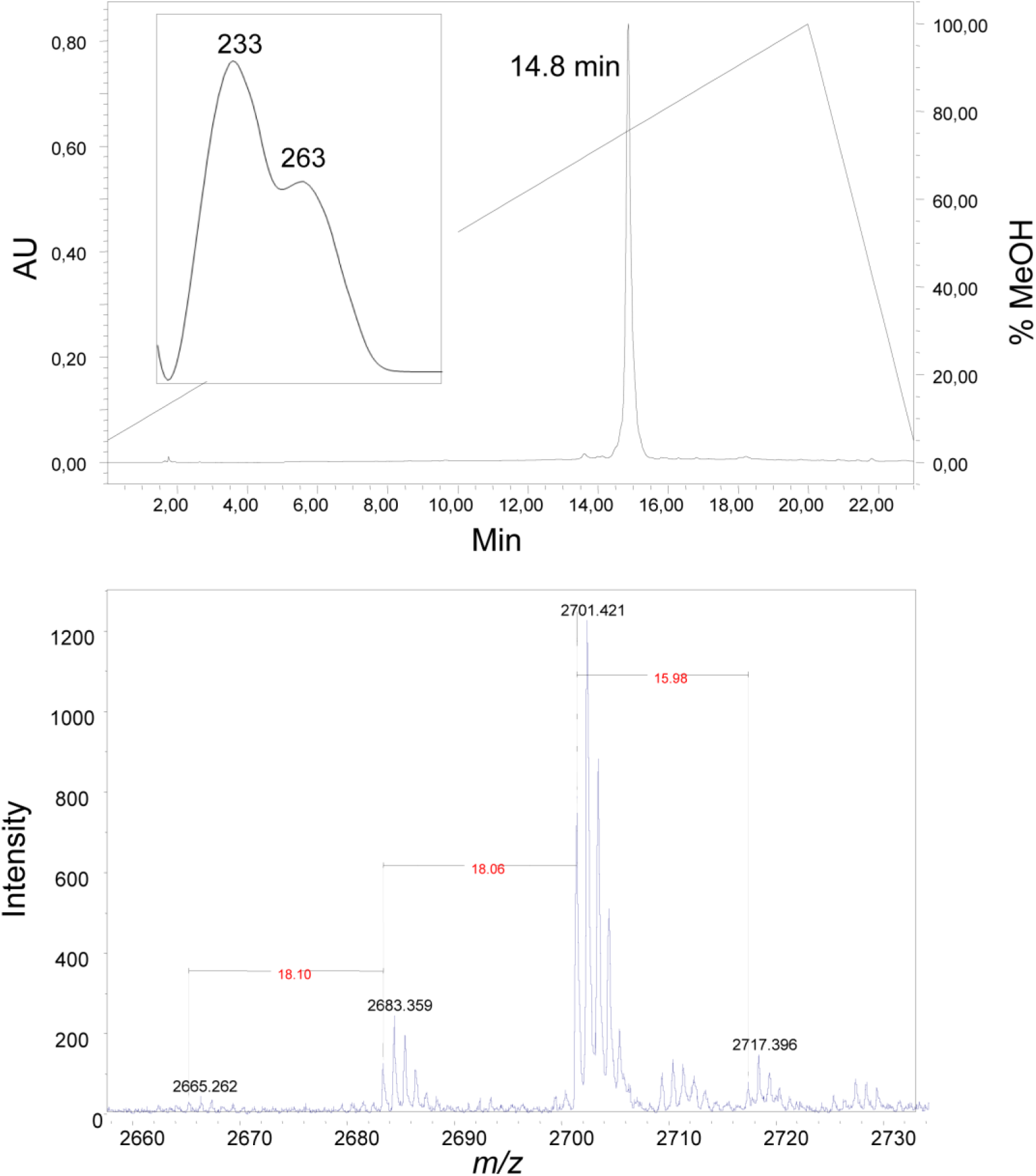
Analysis of palytoxin purified from *Palythoa* sp. Pc001. A. HPLC profile at 263 nm of the palytoxin isolated from *Palythoa* sp. Pc001. The insert graph shows the UV spectrum of the toxin eluted at 14.8 min in the gradient conditions used (see Methods). B. MALDI-ToF mass spectrum of the palytoxin isolated from *Palythoa* sp. Pc001.

The mass spectra corresponded to a single molecule free of contaminants (Fig 3B). Indeed, by using the optimised matrix of Paz *et al*. [48] composed of DHB in a 0.1% TFA/acetonitrile mixture (which gave better resolution than with a α-cyano-4-hydroxycinnamic acid (HCCA) matrix), the MALDI-ToF mass spectrometry in the positive mode revealed single charged molecules containing mainly Na^+^ and few K^+^ ions. The purified toxin showed a clear ion profile with a prominent ion at *m/z* 2701.421 [M + Na]^+^ corresponding to a molecular mass of 2678.43 (Fig 3B). Together with this main molecular ion, minor adducts at *m/z* 2683.359 [M + Na – H_2_O]^+^, 2665.262 [M + Na – 2H_2_O]^+^, and 2717.396 [M + K]^+^ were found, confirming the high degree of PlTX purity. ESI-tandem mass spectrometry analyses on a Q-ToF instrument were also performed to confirm the PlTX structure, and revealed a complex ion pattern with multiply charged states. The cleavage between carbon 8 and 9 of the PlTX was demonstrated by the loss of the A moiety which was recovered as a single fragment ion [M + H]^+^ *m/z* 327.126 in the full MS spectra and corresponded to a C_16_H_27_N_2_O_5_ part formula usually observed in PlTX (data not shown). Other characteristic tri- and bi-charged ions were notably identified at *m/z* 906.495 [M + 2H + K]^3+^, 1351.750 [M + H + Na]^2+^, and 1359.741 [M + H + K]^2+^ (data not show). Further, MS/MS assignations confirmed the presence of a single molecule in the final purified fraction. The calculation based on these tri- and bi-charged ions, together with the MALDI-ToF data, attributed to the PlTX of *Palythoa* sp. Pc001 a molecular weight of 2678.48, similar to that of PlTX of *P. tuberculosa*, *P. toxica*, *P. caribaeorum* and *P. heliodiscus* (Table 1).

### Localisation of PlTX and *Symbiodinium* cells

Cross-sections were analysed by MALDI-IMS to localise the PlTX in *Palythoa* sp. Pc001 tissues. At the same time, chlorophyll and peridinin pigments were analysed by epifluorescence microscopy to localise *Symbiodinium* cells (Fig 4). PlTX was easily identified by MALDI-IMS and displayed a non-homogeneous distribution in polyp tissues. The ectodermal tissues, *e.g.* the epidermis of the body wall and the pharynx, showed the highest concentrations of PlTX (Fig 4A, B). PlTX was also found in some tissues of endodermal origin but with concentrations usually lower than in the ectodermal ones (Fig 4A-F). Amongst the endodermal tissues, the outer side of the endodermal fold close to the epidermis and the inner layer around the pharynx contained the highest PlTX concentrations (Fig 4A, B). Very few PlTX was detected in the gastrodermis of the enteron and septa (Fig 4C-F). High concentrations of PlTX were also detected in the mucus-like secretion surrounding the polyps, especially well visible in the sagittal sections executed along the apico basal axis of the body (data not shown). No PlTX was observed in the tentacles whatever the tissue considered (Fig 4A, B). Sections were also analysed by epifluorescence microscopy to localise the endogenous auto fluorescence due to chlorophyll and peridinin pigments of the *Symbiodinium* cells (Fig 4G-I). A strong signal was detected in the epidermis of both the body wall and the tentacles, the endodermal fold (outer layer of the gastrodermis below the epidermis) at the level of the mouth opening and the inner layer of the gastrodermis constituting the enteron wall. The gastrodermis constituting the septa walls located in the median part of the body also contained numerous *Symbiodinium* cells (Fig 4H), the number of which significantly decreased towards the apex (Fig 4G) and the base (Fig 4I).

**Figure 4.**
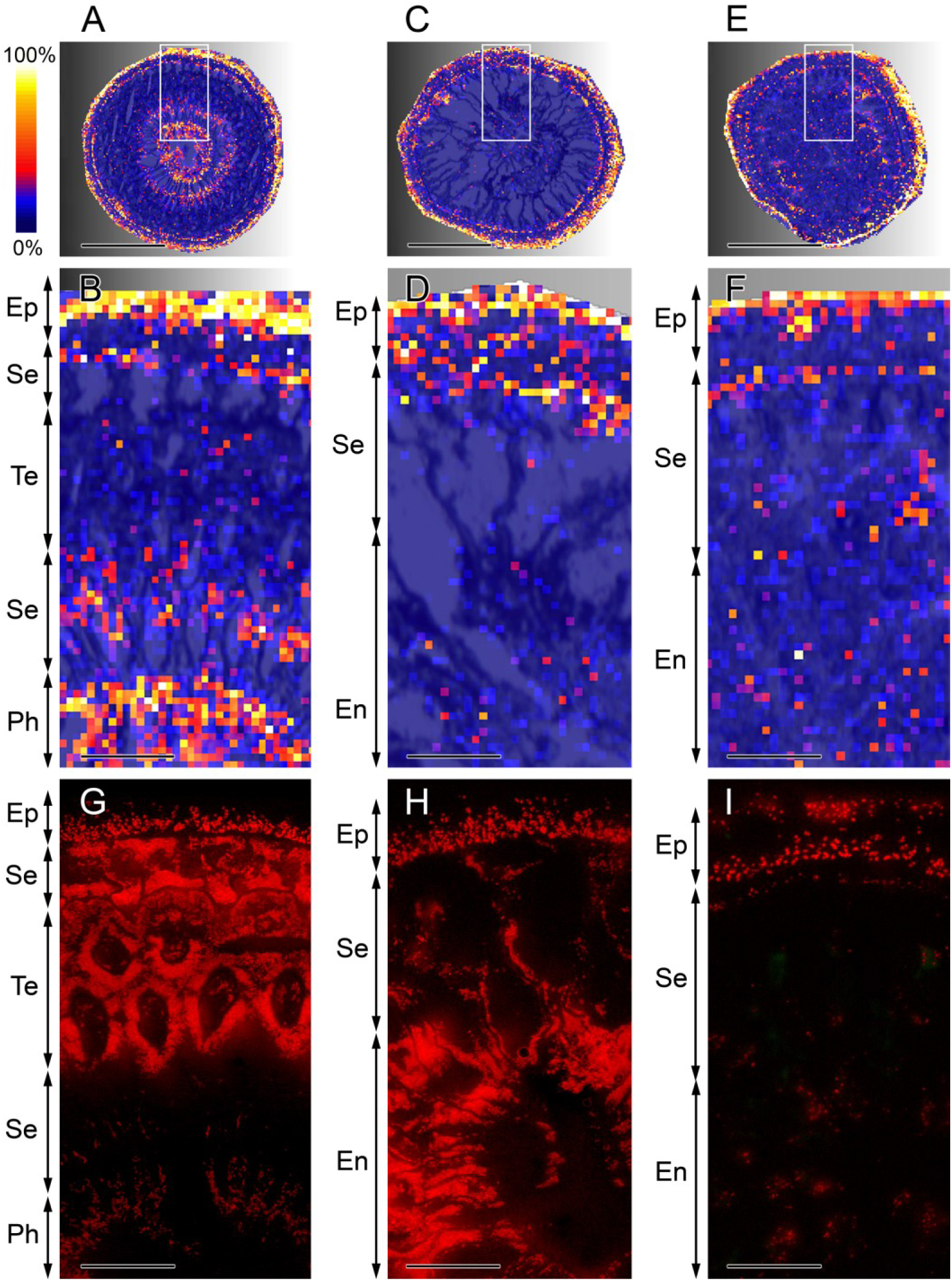
Cross sections of *Palythoa* sp. Pc001. Sections at the mouth and tentacles (A), actinopharynx (C) and basal (E) levels demonstrating the location and relative concentrations of palytoxin (PlTX) by MALDI-imaging mass spectrometry. Areas delimited by white rectangles are enlarged (B, D, F) and compared with images of the same histological regions showing endogenous autofluorescence due to photosynthetic pigments of *Symbiodinium* cells (G-I). Abbreviations: En, enteron; Ep, epidermis; Ph, pharynx; Se, septa; Te, tentacles. Color scale, highest PlTX concentrations (100%) - no PlTX (0%). Scale bars = 3 mm (A, C, E), 0.5 mm (B, D, F, G-I).

### *In vitro* growth inhibitory effect on cancer cells lines and IC_50_

The data obtained by means of the MTT colorimetric assay with PlTX in terms of *in vitro* growth inhibitory effects in human non-cancerous versus rodent and human cancer cell lines were highly reproducible as illustrated in Table 2. Our results revealed that PlTX-mediated sensitivity of the non-cancerous and non-transformed NHDF cell line was ~10^6^ orders of magnitude lower than that of the cancerous or HBL-100 transformed cell lines. Surprisingly, the HBL-100 cell line transformed for immortalisation behaved as a cancer, not as a non-cancerous normal cell line regarding the PlTX growth inhibitory effects.

**Table 2.** IC50 *in vitro* growth inhibitory concentration by 50% after having cultivated the cells in presence of the *Palythoa* sp. Pc001 palytoxin for 72 h.

| Cell line | Phenotype | Origin | $IC_{50}$ (pM) |
| --- | --- | --- | --- |
| NHDF (dermal fibroblast) | Non-cancerous normal cell | Human | $> 1 \times 10^6$ |
| HBL-100 (mammary epithelial) | Non-cancerous immortalised cells | Human | 0.65 |
| A549 (lung carcinoma) | Cancer | Human | 0.67 |
| Hs683 (glioma) | Cancer | Human | 0.58 |
| U373n (glioma) | Cancer | Human | 0.56 |
| 9L (gliosarcoma) | Cancer | Rat | 0.39 |
| B16F10 (melanoma) | Cancer | Mouse | 0.44 |
| Mean ± SEM | Cancer | Human and murine | $0.53 \pm 0.05$ |

For calculation of the average, the $IC_{50}$ value for HBL-100 transformed cell line was included.

The incomplete competition between PlTX and ouabain over the binding of the NaK α1-subunit suggests that this subunit is the main target of PlTX [2]. To test this hypothesis, two rodent models were included in our assay, *i.e.* the B16F10 mouse melanoma and the 9L rat gliosarcoma cell lines. Indeed, rodents display a double mutation in the NaK α1-subunit, they are therefore 100-1,000 times less sensitive to NaK α-subunit inhibitors like the CG UNBS1450, when compared to human cells [55-57]. Interestingly, the rodent cell lines displayed similar sensitivity to PlTX-mediated *in vitro* growth inhibition when compared to human cells (Table 2). The mean IC_50_ calculated for cancer cell lines including the IC_50_ obtained for transformed HBL-100 cells was 0.53 ± 0.05 pM PlTX. Videomicroscopy data illustrate the morphological pictures obtained with 0.01 and 1 nM PlTX on human Hs683 and U373 glioma apoptosis-resistant cells (Fig 5). The concentration of 0.01 nM (*i.e.* 10 pM) represents ~20 times the IC_50_ *in vitro* growth inhibitory concentration associated with PlTX (Table 2), while 1 nM represents ~2,000 times this IC_50_ concentration. Both Hs683 and U373 cell lines began to die about 3 h and 5 h respectively after having been treated with 0.01 nM PlTX. Both cells died after 3 h post-treatment with 1 nM of PlTX. The morphological appearance of dying Hs683 and U373 cells treated with 1 nM PlTX (Fig 5) is typical of cell swelling and bubbling occurring after the impairment of various ion channels and pumps, not only NaK.

**Figure 5.**
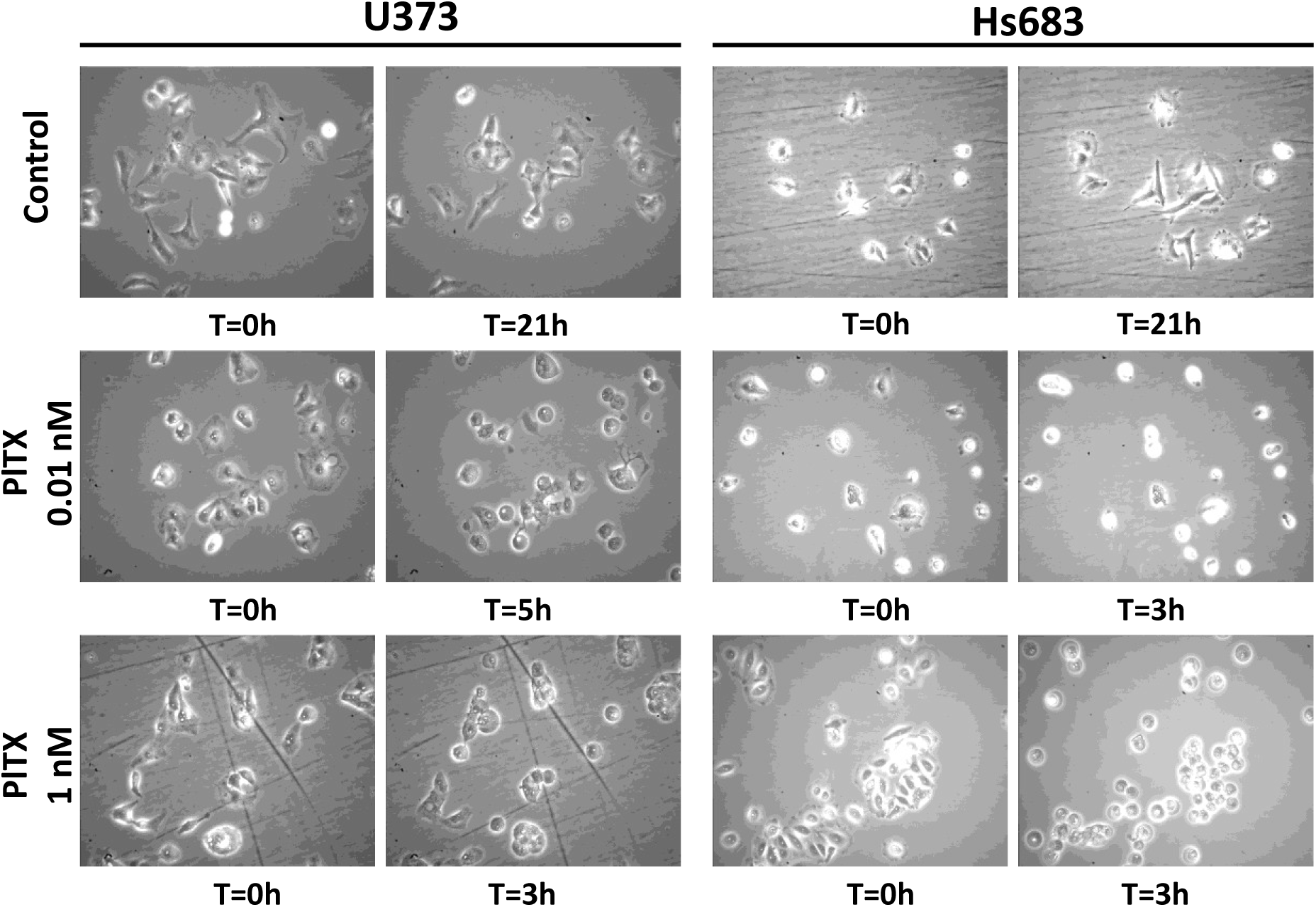
Effect of palytoxin on human cell lines monitored by videomicroscopy. Human U373 (left panel) and Hs683 cell lines (right panel) treated with 0.01 or 1 nM palytoxin. As marked morphological changes of U373 and Hs683 cell lines were observed after 3 to 5 h of treatment with palytoxin, these timepoints were chosen for their illustrations while growth of the control has been followed and is presented at 21 h.

## Discussion

### Taxonomic remarks on *Palythoa* sp. Pc001 and its *Symbiodinium* clade

The identification of the specimens used in this study is a crucial step given that (*i*) different *Palythoa* species might produce different types or stereo-isomers of PlTX, (*ii*) the IC_50_ found here are remarkably lower than previous published data, a result that can be explained by differential pharmacological effects related to different isomers of PlTX [58]. The unambiguous identification of a zoanthid species is not a trivial task, even for an expert taxonomist, mainly due to the lack of reliable diagnostic characters combined with high levels of intraspecific morphological variation [59-61]. Recent molecular studies suggested that several previously described species were simply redundant [61, 62]. Moreover, many of the remaining species exist in sibling pairs between the Atlantic and Indo-Pacific basins. Without collection location or genetic data, it is therefore simply impossible to tell species pairs apart. Our molecular analyses showed that the closest relative to *Palythoa* sp. Pc001 is *Palythoa* aff. *clavata*, an undescribed species found in Florida waters [62]. More recently, Reimer *et al*. [63] identified new specimens from Ascension Island belonging to the *clavata* complex. The morphology of *Palythoa* sp. Pc001 does not match with any Indo-Pacific *Palythoa* species but fits very well with a specimen from the Cape Verde Islands named *Palythoa* sp. 265 [64] which is also genetically very close to *P.* aff. *clavata* [62]. Unfortunately, we could not include *Palythoa* sp. 265 in our concatenated analysis since ITS-rRNA sequences are not available in public databases. Nevertheless, Reimer *et al*. [64] provided its COI sequence, the phylogenetic analysis of which confirmed a close relationship between *Palythoa* sp. 265 from the Cape Verde Islands and *Palythoa* sp. Pc001 (not shown). Molecular analyses have also shown that the undescribed Indo-Pacific species *P.* aff. *sakurajimensis* is genetically close to *Palythoa* sp. Pc001. However, this molecular result is not congruent with morphological data, emphasizing once more that, in zoanthids, genetic similarities do not necessarily reflect morphological ones. Species boundaries are based primarily on morphology and it is generally assumed that morphological variations reflect reproductive isolation with genetic differentiation. However, genetic studies and fertilization trials producing viable larvae suggest that morphologic and genetic distinctions are not always closely linked in corals (for review [65]). So far, even if it is reasonable to consider *Palythoa* sp. Pc001 as an undescribed species on the basis of its morphology and collection location, it is not possible to assess to which extent hybridization with *P.* aff. *sakurajimensis* occurs naturally on the reef. As found in *P.* aff. *clavata* [63] and *Palythoa* sp. 265 [64], *Palythoa* sp. Pc001 contains a ‘generalist’ *Symbiodinium* belonging to clade C. A comparison of published results with the range of pigments synthesised by the *Symbiodinium* sp. found in *Palythoa* sp. Pc001 suggests that these pigments are synthesised only by the *Symbiodinium* cells (see [47, 66, 67]). This, in turns, supports that *Palythoa* sp. Pc001 seem not to be associated with epizoic or endolithic Cyanobacteria, green algae and diatoms, in contrast to other corals [67].

### Storage of PlTX in *Palythoa* sp. Pc001

*Palythoa* sp. Pc001 contains the highest amount of PlTX ever found in a zoanthid (2.22 ± 0.41 mg/g wet zoanthid). The mean recorded value of 0.22% (w/w) PlTX is high compared to the values found in the literature (see Table 1), and corresponds to eight times more than the first value of 0.027% (w/w) recorded by Moore and Sheuer [5] from *P. toxica*. In *P. heliodiscus*, Deeds *et al*. [11] found a maximum of 1164 µg/g wet zoanthid for PlTX corresponding to 0.12% (w/w). The PlTX synthetic pathway and the putative symbiotic organism(s) involved are entirely unknown [6]. Based on structure similarities between PlTX and zooxanthellatoxins, these last being secreted by *Symbiodinium* spp. [7, 8, 68], it has been hypothesised that the symbiotic dinoflagellates in *Palythoa* are responsible for PlTX synthesis. Moreover, several species of free dinoflagellates are able to produce PlTX and analogues [35, 40, 41, 42]. MALDI-IMS carried out on *Palythoa* sp. Pc001 tissues did not highlight an exclusive colocalisation of *Symbiodinium* cells and high levels of PlTX. Strikingly, high levels of PlTX could be detected in histological regions where few or no *Symbiodinium* cells could be observed, notably in the epidermis forming the pharynx. This raises interesting questions about the storage process in toxic *Palythoa* species. PlTX is unlikely to be able to diffuse through cellular membranes due to its large molecular mass. Hence the hypothesis of a central role of the *Symbiodinium* cells in PlTX synthesis suggests that (*i*) PlTX migrates through the symbiosome membranes and the cytoplasmic membrane of the host cell thanks to a non-diffusive active transport and (*ii*) a storage pathway must exist in some cells of toxic *Palythoa* species. It is noteworthy that the PlTX and its analogues are not only produced by dinoflagellates. PlTX and one of its analogues, 42-hydroxy-PlTX, have been shown to be produced by a marine cyanobacteria belonging to the genus *Trichodesmium* [38]. Frolova *et al*. [9] detected PlTX-like compounds in Gram-negative *Aeromonas* sp. and *Vibrio* sp. bacteria using anti-PlTX antibodies. Similarly, bacteria isolated from *Palythoa caribaeorum* were found to display a PlTX-like haemolytic activity [10] confirming that several prokaryotic organisms can produce at least one PlTX type. It is therefore possible to assume that the *Symbiodinium* spp. and symbiotic prokaryotes collaborate with the cnidarian host to produce and store PlTX. Accordingly, the holobiont itself has a decisive role and provides a favourable environment where crucial organisms for the synthesis and storage of PlTX are united. This could also explain the differences of toxicity between several genetically close species of zoanthids [6, 11, 34] containing *Symbiodinium* spp. belonging to the same clade. In any event, identifying the organisms involved in the PlTX synthesis would necessitate both confocal laser scan microscopy and immunolabelling methods associated with metabolomics.

### PlTX from *Palythoa* sp. Pc001 is the most powerful anti-cancer CGs ever reported

Ouabain-related *in vitro* IC_50_ growth inhibitory effects are in the 10-100 nM ranges of concentration [55, 56], thus 10^4^ to 10^5^ orders of magnitude higher than the 0.54 ± 0.05 × 10^−12^ M (picomolar) IC_50_ evaluated in this work against various cancer cell lines. UNBS1450 inhibits α3β1 NaK at 3.2 nM and demonstrates a much higher inhibition of NaK than classic CGs, with potency > 200 times greater than ouabain and ∼6 times greater than digoxin [55]. However, the use of UNBS1450 at antiproliferative concentrations does not induce apoptosis or intracellular Ca^2+^ increases that are typically involved in CG-mediated cardiotoxicity [53]. Videomicroscopy experiments showed that in presence of 10 pM of PlTX isolated from *Palythoa* sp. Pc001, Hs683 oligodendroglioma and U373n glioblastoma cell lines died but with a delay for the pro-apoptotic resistant U373n cells. Both cell lines displayed swelling and bubbling behaviours characteristic to NaK impairment as well as other ion channels. It is known that changes in ion fluxes are the immediate effects of PlTX on the cells. In particular, the increase of the Na^+^ permeability leads to the membrane depolarization and to a secondary Ca^2+^ influx that may lead to multiple events regulated by Ca^2+^-dependent pathways [24, 25]. However, it has been shown at low concentrations (10−100 nM) that other CGs do not affect the ionic imbalance of the cell and bind nonpumping NaK localised in caveolae leading to activation of the Src tyrosine kinase and MAPKs [26, 29]. Several works previously described the effects of PlTX on proliferation and survival of cancer cells in similar experimental conditions, *i.e* without a pre-treatment with ouabain. Valverde *et al*. [69] described the cytotoxic effect triggered by the *P. caribaeorum* PlTX on neuroblastoma cells (ATCC CRL-2267). A 24 h treatment with 1 nM of PlTX inhibited up to over 50% cell proliferation. Significant toxic effects of the *P. caribaeorum* PlTX were also observed in head and neck squamous cell carcinoma cell lines with a LD_50_ of 1.5-3.5 ng PlTX/mL (0.56-1.30 nM), in contrast to healthy epithelial cells [1]. Furthermore, Kerbrat *et al*. [38] reported an IC_50_ of 170 pM with neuroblastoma cells (ATCC CCL-131) incubated for 22 h with a PlTX mixture mainly composed of the 42-hydroxy isomer from the cyanobacteria *Trichodesmium* sp. Finally, Ledreux *et al.* [70] reported an IC_50_ value of 42.9 ± 3.8 pM in experiments carried out on Neuro2a cell line with a PlTX incubation time of 19 h.

What could explain the higher cytotoxicity of the PlTX isolated from *Palythoa* sp. Pc001? The molecular weight of 2679 Da and the fragmentation recorded by mass spectrometry show that the *Palythoa* sp. Pc001 toxin is a ‘classic’ PlTX similar to that isolated from *P. toxica*, *P. tuberculosa and P. caribaeorum* [5, 30, 31]. However, different stereo-isomers may be isolated from different *Palythoa* species. For instance, successive NMR-based stereostructural studies revealed that the two 42-hydroxy-PlTXs isolated from *P. toxica* and *P. tuberculosa* were diastereo-isomers with inverted configurations at C50 [58]. Interestingly, the cytotoxicity of 42-hydroxy-50R-PlTX from *P. tuberculosa* toward skin HaCaT keratinocytes appeared approximately two orders of magnitude lower than that of PlTX and one order of magnitude lower than that of 42-hydroxy-PlTX isolated from *P. toxica* [58]. The configurational inversion at C50 likely causes the 42-hydroxy-50R-PlTX to undergo conformational changes that ultimately reduce its toxicity [2, 58, 71].

### PlTX from *Palythoa* sp. Pc001 acts through unknown NaK binding sites

Ouabain, a potent blocker of the NaK used to inhibit some PlTX effects *in vitro*, can displace PlTX binding on purified NaK [18, 72]. The binding site for ouabain is restricted to the N-terminal 200 amino acids of the NaK α-subunit [73]. Data obtained here suggest that the PlTX binding abilities are not only restricted to binding site for ouabain on the NaK α-subunit. Indeed, we showed similar *in vitro* growth inhibitory activity in rodent and human cancer cells although the NaK in rodents displays double mutation in the α1-subunit which decreases the sensitivity to NaK inhibitors like UNBS1450 from 100 up to 1,000 times, when compared to human cells [55-57]. Furthermore, it has been demonstrated that PlTX and ouabain could simultaneously bind to NaK [74] and a study on immortal human skin keratinocytes (HaCaT) supports the existence of both ouabain-sensitive and -insensitive PlTX binding sites [2]. Together with our results, all of the above strengthens the hypothesis that PlTX-induced cytotoxic effects via NaK might be mediated by binding sites that are distinct and/or partially overlapping with the α-subunit traditionally identified for other CGs.

### PlTX from *Palythoa* sp. Pc001 showed a specific cytotoxicity towards cancer cells

PlTX from *Palythoa* sp. Pc001 shows an exceptionally *in vitro* growth inhibitory activity on cancer cells which is ~10^6^ orders of magnitude higher than on non-cancerous normal cells. This is the broadest differential cytotoxicity of potent NaK ligands ever reported [55-57]. Moreover, the *in vitro* growth inhibitory activity induced by PlTX on cancer cells, associated with sensitivity to pro-apoptotic stimuli, was similar to the activity observed in those associated with various levels of resistance to pro-apoptotic stimuli. CGs are preferentially toxic to melanomas *via* the inhibition of NaK [75]. The CG UNBS1450 is significantly less active (in terms of *in vitro* growth inhibition) in normal fibroblasts (WI-38) than in NSCLC (A549 andCal-12T; [55]) and glioblastoma cells (T98G and U373; [57]). Considering the high cytotoxicity at very low doses towards cancer cells demonstrated here, PlTX from *Palythoa* sp. Pc001 represents a very promising anti-cancer agent. However, due to the lack of knowledge concerning the mechanisms of action of PlTX *via* NaK, it is difficult to explain this puzzling differential cytotoxicity. Several physiological explanations related to different cancer phenotypes can nevertheless be suggested. First, the number of NaK might be different in various cancer cells. While the α1 subunit of NaK is highly expressed in glioblastomas compared with normal tissues [57], Rajasekaran *et al.* [76] showed a down regulation of the β1-subunit in renal cell carcinoma accompanied by a significant reduction in the NaK activity. A low NaK activity may consequently increase the PlTX toxicity in some cancer cells. Second, one can also suggest that the heterodimers constituting the NaK in cancer cells are different and able to increase the death consecutive to the binding of PlTX on particular isoforms. However, several publications emphasised the altered expression of NaK subunits in cancer cells when compared to corresponding normal ones. Different cancer types are characterised by over-expression of different α- and β-subunits and the presence of specific additional regulatory polypeptides, like FXYD proteins or transporters modifying the NaK kinetic properties [56]. Targeting the metabolic differences between cancer and normal cells holds promise as a novel anti-cancer strategy [77]. Cancer cells usually use high level of ATP for their enhanced glycolytic metabolism (Warburg effect, [78]). Distinct UNBS1450-mediated *in vitro* growth inhibitory effects between normal and cancer cells seemed to be correlated with a lower ATP level in cancer than in normal cells [57]. Therefore, cancer cells might compensate less efficiently the ionic imbalance after a PlTX treatment than healthy cells because of a lower ATP level.

Given that, the above observations raise two questions. Does the treatment with PlTX interfere with the glucose metabolism and preferentially reduce ATP level in cancer cells? Is the ATP level in cancer cells sufficient to maintain a healthy metabolism after incubation with picomolar doses of PlTX? To explore that, further experiments are needed. In particular, it will be useful to study intracellular signalling pathways and modification of the ATP level triggered by the PlTX both in cancer and normal cells, in addition to transcriptome analysis using next-generation sequencing in regard to cancer metabolism and PlTX cytotoxicity.

## Authors’ contributions

LS and YP carried out the molecular experiments and phylogenetic analyses. LS and JL did the PlTX purification, HPLC and mass spectrometry experiments as well as pigment identification. LS, CN and JL did the MALDI-IMS analyses. FG maintained the clonal population of *Palythoa* sp. Pc001. JL, YP and LS drafted the manuscript. JL and YP conceived and supervised the study. All authors read, amended and approved the final manuscript.

## Acknowledgements

We are deeply grateful to Céline Bruyère and Véronique Mathieu who provided cell lines and carried out MTT colorimetric assay and quantitative videomicroscopy experiments. Our manuscript benefits from stimulating discussion with Véronique Mathieu and Daniel Papillon who also provided English corrections. This study was financially supported by Coral Biome and Agence Nationale pour la Recherche et la Technologie (France), and IRD. Ludovic Sawelew is working for his PhD project under the CIFRE agreement 2014/0245 (Industrial agreement between Coral Biome and IRD).

